# A non-canonical histone acetyltransferase targets intragenic enhancers and regulates plant architecture

**DOI:** 10.1101/2020.02.25.965475

**Authors:** Xueyong Yang, Jianbin Yan, Zhen Zhang, Tao Lin, Tongxu Xin, Bowen Wang, Shenhao Wang, Jicheng Zhao, Zhonghua Zhang, William J. Lucas, Guohong Li, Sanwen Huang

## Abstract

Axillary meristem development determines both plant architecture and crop yield; this critical process is regulated by the TCP transcription factor (TF) family, including the maize *TB1* and Arabidopsis *BRC1*. Studies have shown that both TB1 and AtBRC1 can target the gene body regions of some target genes and activate their expression; however, the regulatory mechanisms remain largely unknown. Here, we show that a cucumber *CYC/TB1* homologue, *TEN*, controls the identity and mobility of tendrils. Through its C-terminus, TEN binds at intragenic enhancers of target genes; its N-terminal domain functions as a novel, non-canonical histone acetyltransferase (HAT) to preferentially act on lysine 56 and 122, of the histone H3 globular domain. This HAT activity is responsible for chromatin loosening and host gene activation. The N-termini of all tested CYC/TB1-like proteins contain an intrinsically disordered region (IDR), and despite their sequence divergence, they have conserved HAT activity. This study discovered a non-canonical class of HATs, and as well, provides a mechanism by which modification at the H3 globular domain is integrated with the transcription process.

---

TEOSINTE BRANCHED 1 (TB1), CYCLOIDEA (CYC), and PROLIFERATING CELL FACTORS (TCP) transcription factors (TFs) constitute a plant-specific gene family involved in a broad range of developmental processes^1^. Among them, the CYC/TB1 clade of the TCP proteins plays central roles in controlling development of axillary buds that give rise to either flowers or lateral shoots^1,2^. In maize (*Zea mays* L.), the major domestication gene, *TB1*, suppresses branch outgrowth, a crucial architectural modification that transformed teosinte into a viable crop^3^. Subsequent studies on its homologues in rice^4^ and *Arabidopsis thaliana*^5^ identified their similar essential roles in repressing axillary bud growth.

Recently, a genome-wide binding profile uncovered a genetic pathway putatively regulated by TB1^6^. The study reported that TB1 binds mainly to promoters, with only a few peaks located within gene body regions. Nevertheless, other studies have also shown that TB1, and its homologue BRANCHED1 in *Arabidopsis*, can bind to the gene bodies of the target genes *Tassels Replace Upper Ears1* (*Tru1*)^7^ and *HOMEOBOX PROTEIN53*^8^, respectively, to activate their expression. However, the mechanism underlying how the intragenic binding of CYC/TB1-like TFs regulates gene expression is still unclear. Understanding the conserved regulatory mechanism, associated with the function of these CYC/TB1-like proteins, would provide insight into their core role in signal integration of axillary bud repression. Such knowledge could broadly benefit crop breeding programs for tailored plant architecture.

In eukaryotes, enhancers are *cis*-acting DNA sequences which, when bound by specific TFs, increase the transcription in a manner that is independent of their orientation and distance relative to the transcription start site^9^. In *Drosophila melanogaster*, the vast majority (88%) of all enhancers were shown to be located in the vicinity of their targets, of which 30% are upstream, 22% are downstream, and interestingly, 36% are intragenic^10^. These intragenic enhancers appear to mainly (79%) regulate their host genes with only 21% activating neighboring genes^10^. The first eukaryotic intragenic enhancer was discovered in the immunoglobulin heavy chain gene^11^, and it was shown that this enhancer activity was correlated with an increase in histone acetylation and general sensitivity to digestion by DNase I^12^. Several models have been proposed regarding the action of enhancers in the regulation of transcription, of which a ‘facilitated tracking’ mechanism is of interest. In this model, an enhancer-bound complex, containing DNA-binding TFs and coactivators, scans along the chromatin until it encounters the promoter, where a looped chromatin structure is formed. The key points of this tracking mechanism are altering a repressive chromatin structure and facilitating enhancer-promoter communication^9^. Overall, the mechanism regarding transcriptional regulation, by enhancers, is still poorly understood^13^.

The dynamics of chromatin structure are strictly regulated by multiple mechanisms, including post-translational modification of histones^14,15^. Tail-based histone acetylation functions as docking sites for the recruitment of transcriptional regulators, whereas recent data suggest that acetylation of lysine residues, in the globular domain of histone H3 (H3K56 and H3K122), can directly alter histone – DNA interactions, thereby modulating chromatin architecture^16-18^ and stimulating transcription^19-21^. In yeast, histone chaperone-dependent acetylation of H3K56, by the histone acetyltransferase (HAT) Rtt109, is required for chromatin assembly, during DNA replication^22-25^, and for chromatin disassembly during transcriptional activation^19,21^. H3K56 acetylation enhances the unwrapping of DNA, close to the DNA entry–exit site of the nucleosome ^26^, and regulates chromatin at a higher-order level^27^. It also appears that H3K56 acetylation is involved in transcription elongation^28,29^. Similarly, in humans and *D. melanogaster*, the HATs CBP and p300 mediate the acetylation of H3K56, in an Asf1-dependment manner, which is required for chromatin assembly during DNA synthesis^30^. H3K122ac directly affects histone–DNA binding and stimulates transcription^20^. Recently, it was reported that a subset of active enhancers is marked by histone H3 globular domain acetylation (H3K64ac and H3K122ac)^31^. However, the mechanism underlying *cis* and *trans* determinants of how the histone globular domain acetylation is integrated into specific genes, during transcriptional regulation, remains to be elucidated^32^.

Intrinsically disordered regions (IDRs) are polypeptide segments that lack sufficient hydrophobic amino acids to mediate co-operative folding, and thus, lack an ordered three-dimensional structure^33,34^. IDRs are abundant in eukaryotic proteins, being especially prevalent in TFs, and were recently considered to play important roles in gene activation, through the formation of biomolecular condensates (phase separation)^13,35^. However, the possible role and mechanism of action of IDRs, in transcriptional regulation, remains largely unexplored.

Recently, we identified the cucumber (*Cucumis sativus* L.) tendril identity gene, *TEN*, which belongs to the *CYC/TB1* clade of the *TCP* gene family^36^. Tendrils are modified branches in which axillary meristems are inhibited from developing, and climbing behavior is acquired. To understand how TEN regulates target gene expression, we investigated the genome-wide binding profiles of TEN. We show that TEN acts both as an intragenic enhancer-binding TF and as a novel, non-canonical HAT, acting on H3K56 and K122 for host gene activation. Furthermore, we demonstrate that the N-termini of tested CYC/TB1-like proteins contain intrinsically disordered regions (IDRs), and despite their sequence divergence, they have conserved HAT activity.

## Results

### Regulation of tendril identity requires TEN N- and C-termini

A cucumber *TEN* mutant forms modified branches, instead of tendrils, and had therefore lost the capacity to climb^36^. This *ten* gene had a single-point mutation (asparagine to tyrosine at the 338^th^ amino acid residue; N338Y; *ten-1* mutant) in the TEN C-terminus, indicating an important function associated with this region^36^ (Fig. 1a and Supplementary Fig. 1a-b). In order to knock out the *TEN* gene, and further explore TEN-associated functions, we employed CRISPR-Cas9 to target the TEN N121 region (amino acids 1 to 121 in the N-terminus) (Fig. 1a). These *TEN*-edited plants were phenotyped, and a null-mutant (*ten-2*; Fig. 1b) displayed a complete transformation of its tendrils into lateral branches (Fig. 1c-d), equivalent to the *ten-1* mutant phenotype (Supplementary Fig. 1b). This result confirmed the function of TEN in control of tendril identity.

**Figure 1.**
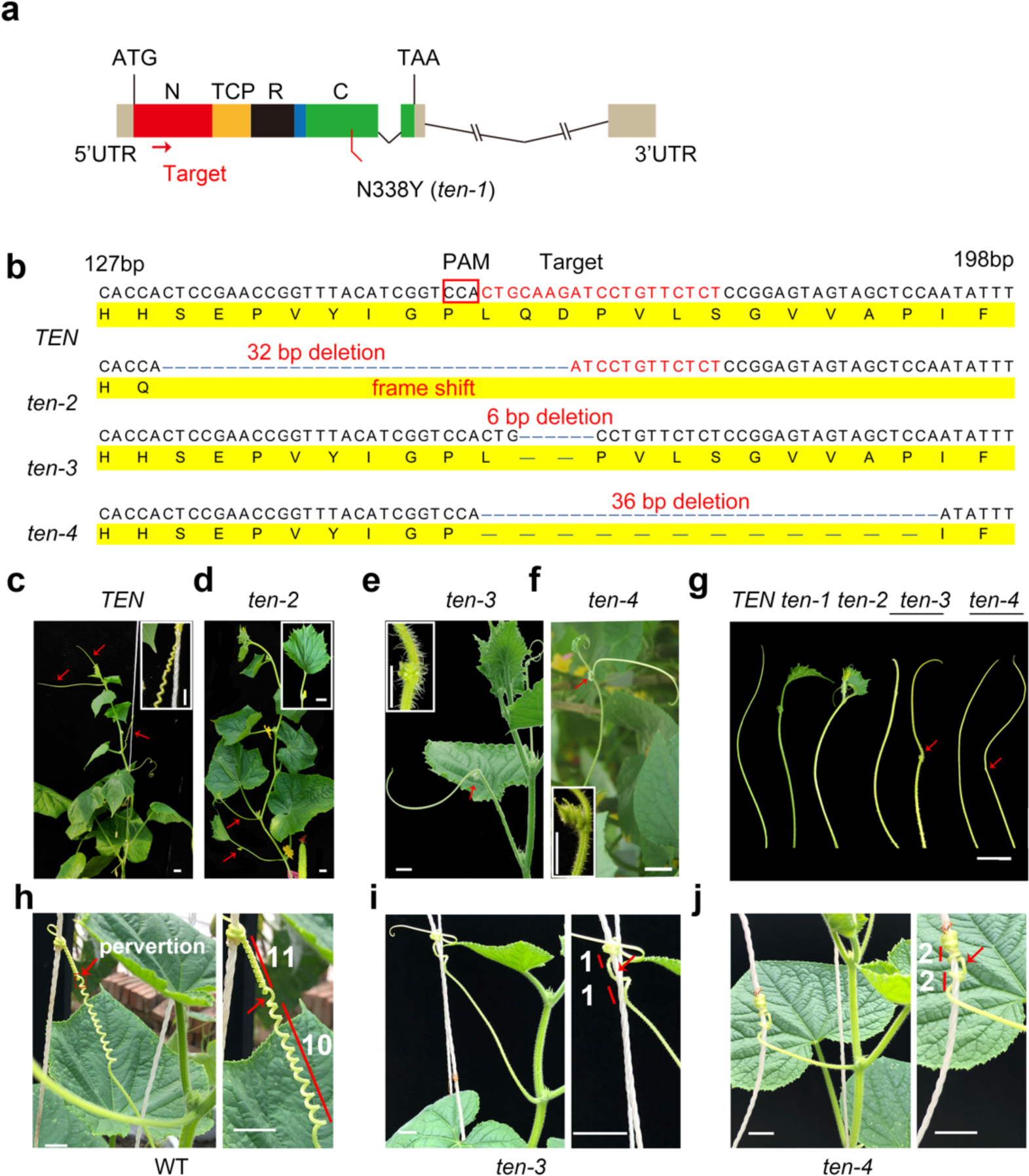
Analysis of CRISPR-Cas9 mutants reveals the *in vivo* role of TEN. **a**, Schematic illustrating the sgRNAs (red arrow) targeting the region in the N121 domain. Colored boxes represent exons, and black lines represent introns. N338Y designates the *ten-1* mutant that forms modified tendrils. **b**, Identification of the *ten-2* allele as a null mutation carrying a 32 bp deletion, and alleles homozygous for genes encoding proteins with small deletions of two amino acids (*ten-3*) and 12 amino acids (*ten-4*). Red font highlights sgRNA targets, and red box indicates the protospacer-adjacent motif (PAM) sequence. **c** and **d**, Wild-type (WT) plant bearing typical tendrils (red arrows), has the ability to climb (**c**), whereas the *ten-2* bears modified tendrils (red arrows) and an inability to climb (**d**). **e** and **f**, Examples of *ten-3* (**e**) and *ten-4* (**f**) tendrils on which axillary buds have developed. **g**, Tendril phenotypes of various alleles for the *TEN* gene. **h**-**j**, Compared to WT (**h**), free coiling, formed by two oppositely handed helices, is impaired in *ten-3* (**i**) and *ten-4* (**j**) plants. The number of turns to each side of the perversion point (shown in white font) indicates the degree of coiling. Arrows indicate the perversion of coiled tendrils. The *ten-3* and *ten-4* displayed a substantial reduction in helical turns on both sides of the perversion points. Scale bars, 2 cm.

In addition to the *ten-2* null-mutant, we also identified two other TEN-edited plants, *ten-3* with a homozygous in-frame deletion of two amino acids (Gln^54^ and Asp^55^), and *ten-4* with a homozygous in-frame deletion of 12 amino acids (Fig. 1b). We observed that in both *ten-3* and *ten-4* plants (Fig. 1e-g), some tendrils retained much of the normal tendril morphology (Fig. 1g); however, interestingly, some showed slight morphological changes, producing axillary meristems on their tendrils (Fig. 1e-g), demonstrating the important role of TEN in axillary meristem inhibition, during tendril development.

Although tendril morphology was largely unaffected in *ten-3* and *ten-4* plants, their climbing capacity was significantly altered (Fig. 1h-j and Movie. 1). Wild-type tendrils form approximately 10 helical turns on each side of a perversion point^37^ (Fig. 1h and Movie. 1). Although these mutant tendrils could still attach to a support, the free coiling, formed by two oppositely-handed helices, was impaired, resulting in a reduced number of helical turns on both sides of the perversion point (Fig. 1i-j and Movie.1). These results demonstrate that TEN controls tendril identity and climbing ability, and that both the N- and C-terminus are critical for its function.

### TEN C-terminus binds at intragenic regions of target genes

To understand how the N- and C-termini of TEN affect TF functions, we first assessed its global binding profiles by chromatin immunoprecipitation sequencing (ChIP-seq). To this end, polyclonal antibodies were raised, and antibody specificity was confirmed (Supplementary Fig. 2a-c and Supplementary Table 1). We also performed immunoblot assays and confirmed that this antibody recognized recombinant full length TEN, but not its TCP domain (Supplementary Fig. 2d). This finding excludes the possibility that this antibody selects against TEN proteins which bind to the DNA via their bHLH domain. ChIP-seq assays were performed using tendrils at the coiling stage. Two replicate experiments were performed and shared a large number of peaks covering more than 59% of peaks in the smaller replicate (Fig. 2a and Supplementary Table 2). Follow-up analysis of genome distribution, using the overlapping peaks, revealed that these TEN-binding sites were highly enriched in intragenic regions (1257 peaks), which accounted for ∼74% of all peaks (65.4% in coding and 8.3% in intronic regions) (Fig. 2b-c). In addition to intragenic regions, a minor portion of the binding sites was distributed in intergenic regions (15.3%), with only 6.1% in promoter regions located within 5 kb upstream of the transcription start site.

**Figure 2.**
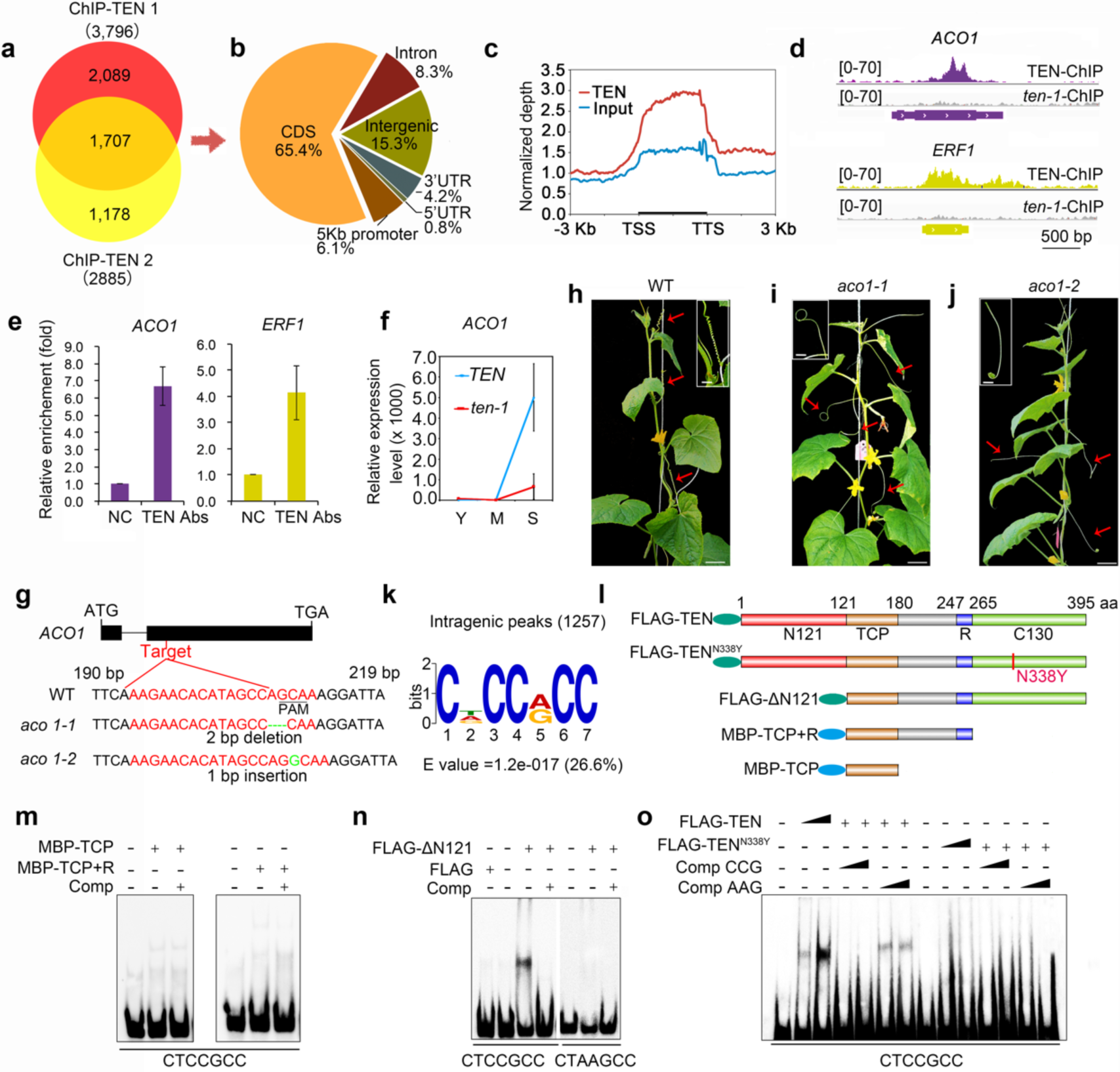
TEN is a novel TF with intragenic binding capacity. **a**, Overlap of TEN binding sites in two TEN ChIP-seq replicates. **b**, Distribution of overlapped TEN binding peaks in the cucumber genome. **c**, TEN binding peaks are highly enriched in the intragenic regions of coiling tendrils. TSS and TTS, transcription start and termination sites. **d**, Two examples of TEN binding profiles in the gene bodies of *ACO1* and *ERF1*. **e**, qPCR analysis of TEN recruitment to the indicated intragenic region (mean ± SEM, *n* = 3). NC, negative control. **f**, Relative mRNA expression levels of *ACO1* during tendril growth detected by RT-qPCR (mean ± SEM, *n* = 3). Y, young; M, medium; S, stretch. **g**, Identification of the *aco1-1* and *aco1-2* alleles as two independent null mutations. Red font highlights sgRNA targets, and underline indicates protospacer-adjacent motif (PAM) sequence. **h**-**j**, Compared to wild type (WT) (**h**), the tendrils in *aco-1* (**i**) and *aco-2* (**j**) form irregular coiling and could not attach to their supports. Arrows indicate the coiled tendrils and insets illustrate the differences in coiling between WT and mutants. Scale bar, 5 cm. **k**, The enriched motif CDCCRCC. **l**, Schematics of WT TEN, the N338Y mutant and three truncated proteins. **m**, EMSA showing that TCP and TCP+R do not bind to a DNA probe containing the CTCCGCC motif. **n**, FLAG-ΔN specifically binds to DNA containing the CTCCGCC motif. **o**, FLAG-TEN specifically binds to DNA containing the CTCCGCC motif, but FLAG-TEN^N338Y^ does not. Comp CCG, competitor (unlabeled CTCCGCC probe); Comp AAG, mutant competitor (unlabeled CTAAGCC probe); +/-, presence/absence of protein or competitor; closed triangle, increasing amount of protein (1 or 4 μg) or competitor (100- or 1000-times that of labeled probe).

To investigate the regulatory spectrum of TEN, a total of 637 genes associated with 1707 peaks were identified, most of which (474 genes; 74.4%) are intragenic targets, with only 7.2% (46 genes) being putatively regulating in promoter regions (Supplementary Table 3). Therefore, in this study, the 474 genes associated with 1257 intragenic binding sites were designated as the TEN target gene set (Supplementary Fig. 2e and Supplementary Table 4). Gene ontology analysis indicated that some genes, involved in axillary bud formation, such as the *SPL* TF genes^38^ and homeobox-related TF genes^39^, were also present in the target gene set, consistent with the role of TEN in tendril morphology regulation (Supplementary Table 4). Genes involved in ethylene biosynthesis and signal transduction were significantly enriched (*P* < 0.05) (Supplementary Fig. 2f). Exogenous spraying with ethephon, a plant growth regulator that is converted to ethylene in the plant, induced spontaneous tendril coiling (Supplementary Fig. 2g), consistent with a central role for ethylene in tendril coiling^40^.

Next, we focused on two exemplary target loci, *ACO1* (*Csa6G160180*), encoding a 1-aminocyclopropane-1-carboxylate-oxidase enzyme for ethylene synthesis, and *ERF1* (*Csa7G049230*), encoding an ethylene response factor (Fig. 2d). As revealed by ChIP-quantitative polymerase chain reaction (ChIP-qPCR), TEN was recruited to the exons of both *ACO1* and *ERF1* (Fig. 2e). To provide further genetic evidence that these intragenic targets are genes directly regulated by TEN, we selected the *ACO1* gene for in-depth investigation. *ACO1* is preferentially expressed in tendril tissue (Supplementary Fig. 2h), and its pattern during tendril growth showed that it was upregulated more than 4,000-fold, from the young stage to stretch stage. By contrast, the activation of *ACO1* was repressed, significantly, in the *ten-1* mutant (Fig. 2f), suggesting an important role for *ACO1* in the tendril coiling process. We also found that, from the young stage to stretch stage, although the expression levels of *TEN* were only slightly up-regulated, the TEN’s binding levels on the *ACO1* locus were significantly up-regulated by approx. 4-fold; this pattern is correlated with the expression pattern of *ACO1* (Supplementary Fig. 2i).

To further explore the function of *ACO1*, CRISPR-Cas9 was employed to target the *ACO1* N-terminus, and we identified two null-mutants, *aco1-1* and *aco1-2* (Fig. 2g). We observed that although tendril morphology was largely unaffected in *aco1-1* and *aco1-2* plants, their climbing capacity was altered significantly (Fig. 2h-j and Movie 2). Wild-type tendrils could attach to the climbing supports and showed normal free coiling activity (Fig. 2h and Movie 2); however, these mutant tendrils displayed irregular coiling and could not attach to their supports (Fig. 2i-j and Movie 2). These results demonstrated that *ACO1* is an authentic direct target of TEN, providing genetic evidence that TEN directly regulates its intragenic targets.

To further analyze the summits of intragenic peaks, we identified a statistically overrepresented motif, CDCCRCC (Fig. 2k and Supplementary Fig. 2j). We expressed and purified a series of truncated TEN proteins, using Sf9 insect cells, to investigate TEN binding activity on this motif (Fig. 2l and Supplementary Fig. 2k-m). Electrophoretic mobility-shift assays (EMSAs) and Surface plasmon resonance (SPR) established that the TCP and R domains bind the previously described TCP binding motif GGTCCC, with high affinity, but had no significant affinity for a 50 bp probe containing the CTCCGCC motif (Fig. 2m and Supplementary Fig. 3a-d). However, purified ΔN121 (containing the TCP+R+C130 domains) and full length TEN protein bound to the DNA fragment containing this newly identified CTCCGCC motif, supporting an essential role of the C130 region (amino acids 265 to 395 in the C-terminus) in the sequence-specific DNA binding of TEN (Fig. 2n and Supplementary Fig. 3e-g).

Our EMSA assays performed with full length TEN established that it binds both the previous GGTCCC motif and the new CTCCGCC motif; however, the Kd is lower for the new motif, reflecting stronger binding to this new motif (Supplementary Fig. 3f-g). In addition, we also showed that the TEN protein could bind, specifically, to the CTCCGCC motif, which could not be competed with the GGTCCC probe (Supplementary Fig. 3h). Furthermore, although the N338Y mutation, in the C-terminus of TEN, had no effect on TEN’s binding to GGTCCC motif (Supplementary Fig. 3i), abolished its binding to this CTCCGCC motif (Fig. 2o), which coincided with the phenotypic changes induced by the N338Y mutation in *ten-1*^36^. Lastly, we established that the purified MBP-C protein could not bind to CTCCGCC probes, indicating that, despite the essential role of the TEN’s C-terminus, in binding the CTCCGCC motif, the C-terminus alone is not sufficient for this binding capacity (Supplementary Fig. 3j-k). These findings provided strong support for the notion that TEN is a TF with intragenic binding capacity.

### TEN binding site is a novel intragenic enhancer for host gene activation

To explore the role of TEN in regulating expression of intragenic target genes, the transcriptomes of candidate genes were analyzed in wild-type (WT) versus *ten-1* mutant plants. Among these 474 genes, 132 were downregulated significantly in mutant tendrils (>1.5-fold change; *P*<0.05); interestingly, no gene was upregulated significantly in the mutant (<0.67-fold change; *P* < 0.05; Supplementary Table 4). This result suggests that the intragenic binding sites (CDCCRCC) of TEN appear to have an enhancer activity for its host genes.

To explore whether the binding of TEN, to the potential intragenic enhancer (CDCCRCC), can regulate the neighboring genes, we investigated the expression of genes flanking the TEN intragenic targets (three genes upstream and downstream) in WT and the *ten-1* mutant. These experiments showed that TEN activation occurred predominantly on its specific target genes (Fig. 3a). To validate these data, qRT-PCR was performed on two exemplary targets, *ACO1* and *ERF1*, and their flanking genes, in tendrils of WT, *ten-1, ten-2* and *ten-3* plants. Our results showed that dysfunction of TEN, in these mutants, leads to a significant reduction in both *ACO1* and *ERF1* expression (Fig. 3b and Supplementary Fig. 4a), whereas there was no significant effects on the expression of the flanking genes, placed either upstream or downstream of the *ACO1* and *ERF1* loci (Fig. 3b and Supplementary Fig. 4a).

**Figure 3.**
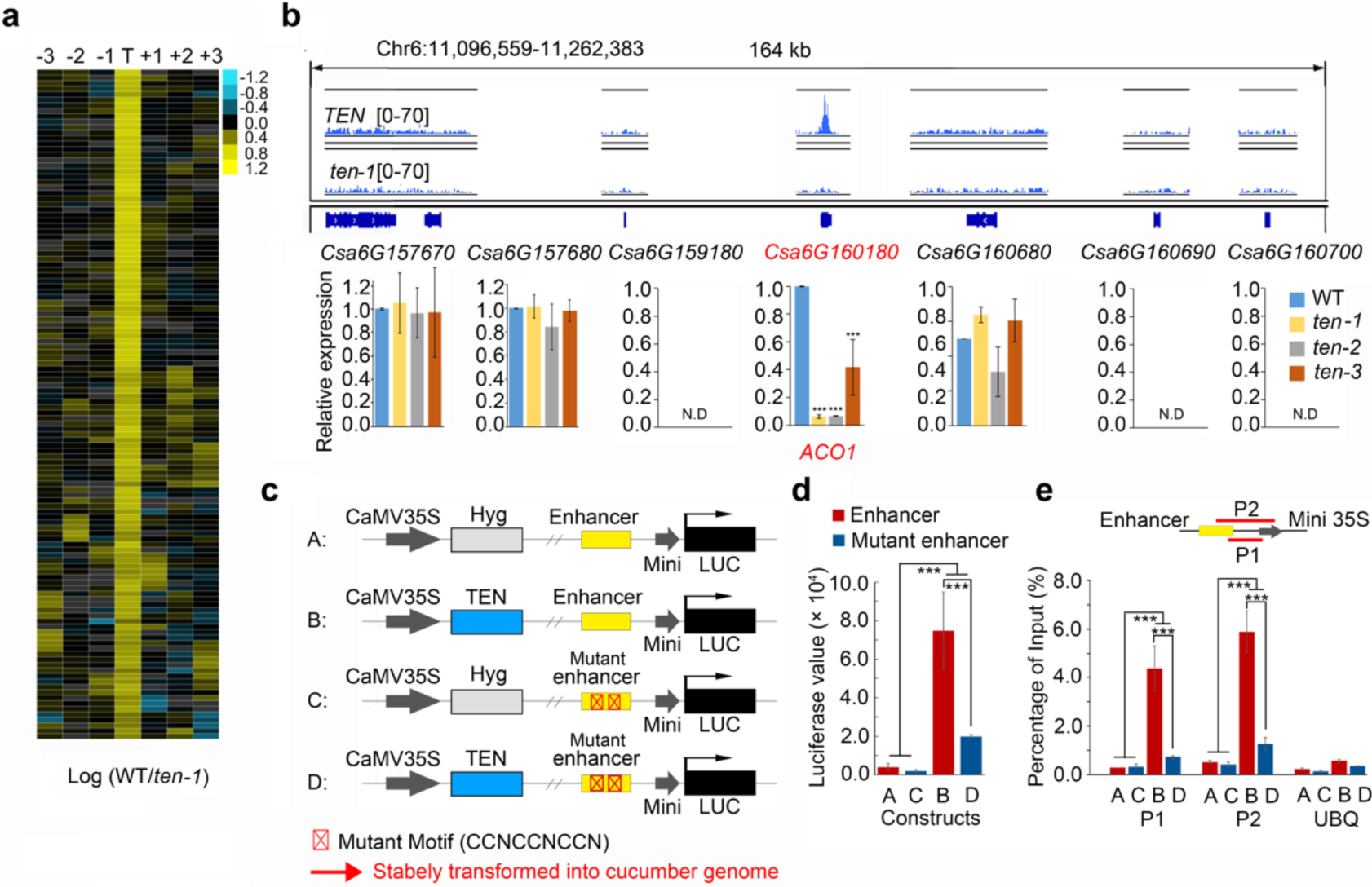
TEN binds to intragenic enhancers of its target genes. **a**, Expression ratio of 132 direct upregulated genes of TEN, between WT and *ten-1*, showing TEN regulates predominantly its host genes. T, intragenic target genes; -1 -2 -3, three genes locating upstream of target gene; +1 +2 +3, three genes locating downstream of target gene. **b**, Expression of putative enhancer target gene, *ACO1*, and flanking genes, assayed by RT-qPCR (mean ± SEM, *n* = 3). *UBQ* was used as internal control. N.D, not detected. **c**, Schematic showing the construction strategy for *in vivo* enhancer validation, through reporter transgenic lines. **d**, LUC activity of positive transgenic cucumber leaves (mean ± SEM, *n* = 4). Four independent transgenic plants, per construct, were used for detection. **e**, ChIP-qPCR analysis of TEN recruitment to the indicated regions (P1 and P2) of the intragenic enhancer (mean ± SEM, *n* = 3). *UBQ* was used as internal control.

To demonstrate, *in vivo*, that the intragenic regulatory elements, bound by TEN, are a novel type of enhancer sequences, and to test whether the CDCCRCC motifs are required for the observed enhancer activity, reporter transgenic lines were assays^41^. To this end, we selected *ACO1* full length genomic DNA sequence, as an enhancer candidate, and each construct contained an expression cassette with TEN, or a hygromycin (*Hyg*) gene under the control of the CaMV *35S* promoter, and another expression cassette containing the enhancer candidate (or mutant enhancer in which we disrupted the respective motifs by point mutations), minimal promoter and luciferase (*LUC*) reporter gene (Fig. 3c and Supplementary Fig. 4b). All four constructs were integrated, independently, into the cucumber genome. Importantly, construct B (TEN + enhancer) exhibited 20-fold higher LUC activity than construct A, the negative control (Hyg + enhancer), and moreover, the construct D (TEN + mutated enhancer) had strongly reduced LUC activity, compared to construct B (TEN + enhancer) (Fig. 3d).

To further confirm that the transformed TEN protein regulates the intragenic enhancer activity, through binding directly and specifically to the intragenic enhancer, we investigated the binding capacity, by ChIP-qPCR, using leaves from plants expressing one of the stably integrated constructs. Here, we demonstrated that TEN could bind the intragenic enhancer, *in vivo*, and that mutation in the CTCCNCCN motif largely impaired this TEN-binding capacity (Fig. 3e). These findings demonstrate that TEN binds on a novel type of intragenic enhancer, and further, validate the functional importance of the CDCCRCC motifs in the enhancer.

### TEN is a new, non-canonical HAT

Having demonstrated that TEN binds to intragenic enhancers of host genes, via its C-terminus, we next explored the functional role of its N-terminus. PSI-BLAST (Position-Specific Iterative Basic Local Alignment Search Tool) analysis revealed that the N121 shares a modest similarity with the transferase domain of an *Arabidopsis* HXXXD acyltransferase (At1G03940; 26% identity, Supplementary Fig. 5a). Considering that the intragenic enhancer-binding TFs might participate in shaping chromatin structure^42^, we speculated that TEN has HAT activity.

To assess this notion, we tested the activity of recombinant TEN protein and determined that it acetylated all four core histones, when histone H3 or free core histones were used as substrates (Fig. 4a). Histone tetramers or octamers enhanced the acetylation ability of TEN, which occurred predominantly in histone H3 (Fig. 4a). TEN also efficiently acetylated mononucleosomes assembled on the 208 bp 5S rDNA, with a preference for nucleosomal histone H3 (Fig. 4b).

**Figure 4.**
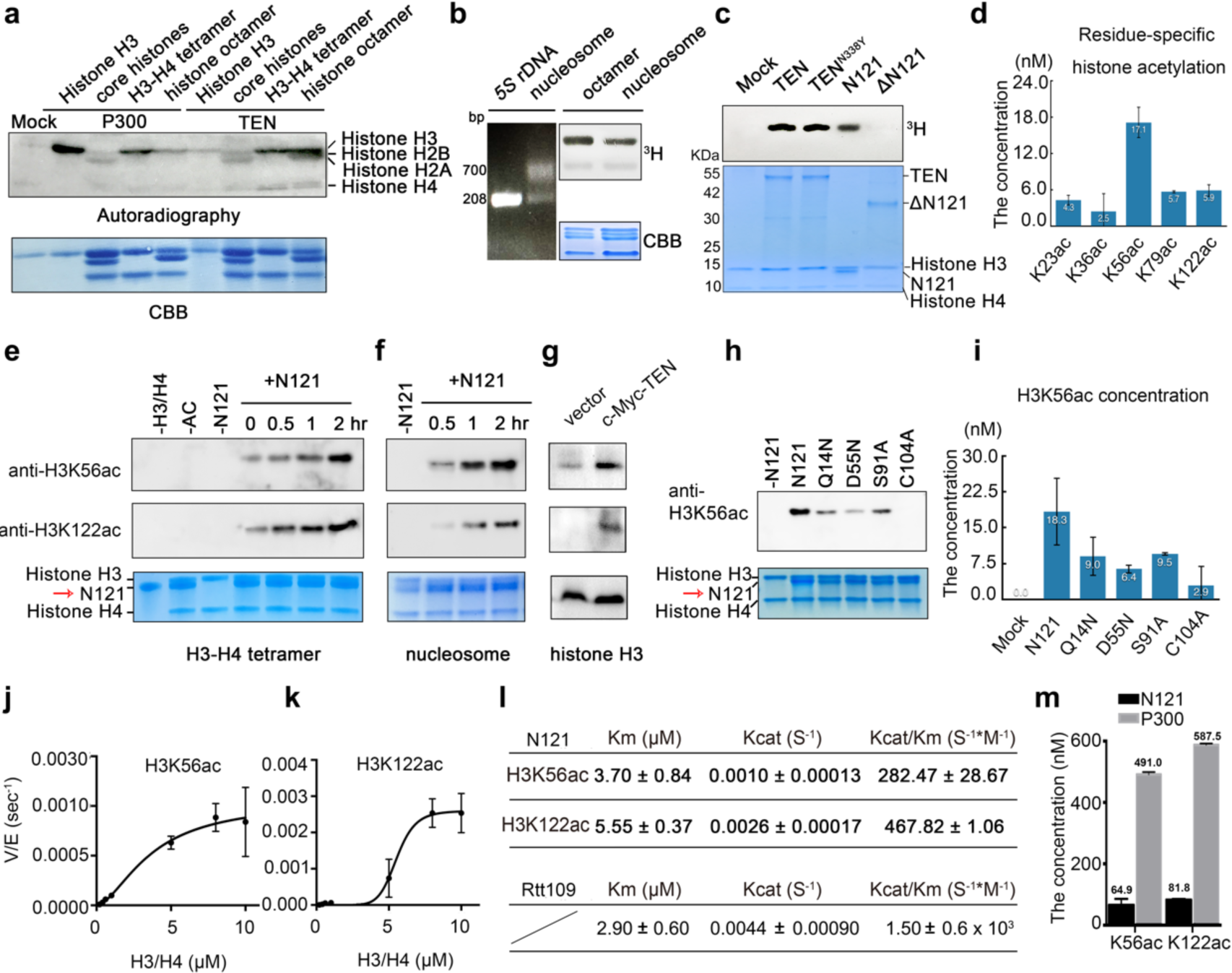
TEN is a novel histone acetyltransferase (HAT). **a**, HAT assays with radiolabeled acetyl CoA and recombinant P300, FLAG-TEN or mock (control). CBB, Coomassie brilliant blue. **b**, Mononucleosomes (∼700 bp) assembled on *5S* rDNA (208 bp), visualized in ethidium bromide-stained gels (left), subjected to HAT assays with recombinant FLAG-TEN with ^3^H-labeled acetyl CoA and visualized by autoradiography or CBB staining (right). **c**, HAT assays with radiolabeled acetyl CoA and FLAG-TEN, FLAG-TEN^N338Y^, FLAG-N121 and FLAG-ΔN121 or mock (control). **d**, Quantification of acetylation levels of individual lysines on histone H3 after the HAT assay using the recombinant N121 protein expressed in and purified from *E. coli*, based on mass spectrometry (mean ± SEM, n=3). **e** and **f**, Acetylation of H3K56 and K122 within the H3-H4 tetramer (**e**) and nucleosome (**f**) by N121 purified from *E. coli*, determined by immunoblotting analysis. **g**, Overexpression of c-Myc-TEN in tobacco leaves increased the acetylation levels of H3K56 and H3K122. **h** and **i**, N121 mutant proteins purified from *E. coli* showed loss of acetylation on H3K56. **j** to **l**, Steady-state kinetics of HAT activity of N121 purified from *E. coli* on H3K56 (**j**) and H3K122 (**k**) (*n* = 3). Derived kinetic parameters for K_m_ and K_cat_ are shown, as compared with Rtt109^42^ (**l**). **m**, A direct comparison between N121 HAT activity and that of a canonical HAT P300, by quantitative mass spectrometry. HAT assays with 600 μM acetyl CoA, 0.2 μM N121/P300, and 2.0 μM H3-H4 tetramer at 30°C for 3 hr. The y-axis indicates the calculated concentration of acetylation at specific lysine residues.

In order to identify the intrinsic HAT domain in TEN, we performed HAT activity assays using recombinant TEN, TEN^N338Y^, N121 and ΔN121 (Supplementary Fig. 5b). The first three forms acetylated histone H3 within H3-H4 tetramers, whereas ΔN121 had no detectable HAT activity (Fig. 4c). The TEN acetylation site specificity was next assessed, by quantitative mass spectrometry^43^, to measure the acetylation levels of individual lysine residues on histones, using the recombinant N121 protein expressed in and purified from *Escherichia coli*. Our *in vitro* assays indicated that N121 acetylated K23 and K36, in the tail domain of histone H3, as well as K56, K79 and K122 in its globular domain, with a preference for K56, K79 and K122 (Fig. 4d and Supplementary Fig. 5c-e).

To confirm these LC-MS/MS results, we performed a combination of *in vitro* HAT assays and immunoblotting, with antibodies specific for different acetylated sites. These assays confirmed that full-length TEN and N121 both acetylated the identified lysines of histone H3, with a preference for H3K56 and K122 (Fig. 4e-f and Supplementary Fig. 5f-g). In addition, the TEN *in vivo* acetylation patterns of H3 lysine residues were assessed by transient expression of exogenous TEN in *Nicotiana tabacum* leaves; here, we observed an increase in acetylation of H3K56 and H3K122 (Fig. 4g and Supplementary Fig. 5h). Furthermore, TEN overexpression led to a significant increase in nuclear H3K56ac and H3K122ac levels, as revealed by immunolabeling of *N. tabacum*, indicating that TEN acetylates, *in vivo*, chromatin-bound H3K56 and K122 (Supplementary Fig. 5i-j). Collectively, these *in vitro* and *in vivo* assays established that the N121 contains the intrinsic HAT domain of TEN, with preferential acetylation of H3K56 and K122 within nucleosomes.

The N121 shares no sequence homology with any other known HAT. To further study its enzymatic mechanism, we therefore focused on some specific amino acid residues, including glutamine (Gln^14^), aspartate (Asp^55^), serine (Ser^91^) and cysteine (Cys^104^) that generally form part of the catalytic center. Point mutations of these candidate residues (Q14N, D55N, S91A and C104A) (Fig. 4h and Supplementary Fig. 6) were tested, using *in vitro* HAT assays, to assess their effects on H3 K56 acetylation. All mutations showed a varying degree of reduction in HAT activity. In particular, the C104A mutant protein had almost no ability to acetylate H3K56 (Fig. 4h-i). These findings demonstrate that C104 is essential for HAT activity, while D55, Q14 and S91 also contribute to HAT activity. The critical role of Asp^55^ for this N121 HAT activity suggested a biochemical basis underlying the phenotypic changes induced by the Asp^55^ deletion in the *ten-3* and *ten-4* mutants.

To gain biochemical evidence that N121 is a *bona fide* HAT, we next conducted steady-state kinetics of H3K56 and K122 acetylation^43^. Here, acetylation of both H3K56 and K122 showed saturation kinetics (Fig. 4j-k). The lower K_m_ for H3K56ac indicated a higher affinity of N121 for H3K56, whereas the catalytic efficiency of H3K56ac was ∼60% of that for H3K122ac (Fig. 4l). We compared the acetylation kinetics of N121 and Rtt109, a HAT required for H3K56 acetylation in yeast. Based on a previous study, we established that both N121 and Rtt109 exhibited similar levels of activity on the H3-H4 tetramer^44^ (Fig. 4l).

We also directly compared the HAT activity of N121 with a canonical HAT P300, by quantitative mass spectrometry. These results showed that N121 acts preferentially on lysine 56 and 122 of histone H3 (Fig. 4m and Supplementary Fig. 7a). The HAT activity of the TEN-N121 on H3K56 and H3K122 was approx. seven times less than that of the canonical HAT P300 (Fig. 4m and Supplementary Fig. 7a).

Taken together, these findings demonstrate that the N121 domain of TEN is a novel histone acetyltransferase which functions, preferentially, at the globular domain of histone H3 within the nucleosome.

### *In vivo* evidence that TEN facilitates chromatin accessibility

To further probe the relationship between TEN binding and acetylation of H3K56 and H3K122, *in vivo*, we mapped H3K56ac and H3K122ac levels, genome-wide, by ChIP-seq and compared the results to the distribution of TEN binding sites of tendril tissue at the coiling stage. The vast majority (>99%) of H3K56ac and H3K122ac peaks overlapped (Fig. 5a). The intragenic binding sites of TEN were colocalized predominantly with the H3K56ac (1,073 out of 1,257 peaks) and H3K122ac peaks (1,054 out of 1,257 peaks, Fig. 5a). In addition, we observed that there were 1,043 TEN binding peaks colocalized with both H3K56ac and H3K122ac peaks (Supplementary Fig. 7b and Supplementary Table 5). H3K56ac and H3K122ac were observed to more likely occur on exons and introns than promoters and intergenic regions bound by TEN (Fig. 5b). These results indicate that the binding of TEN to gene bodies is related to acetylation of the histone H3 globular domain (Fig. 5b).

**Figure 5.**
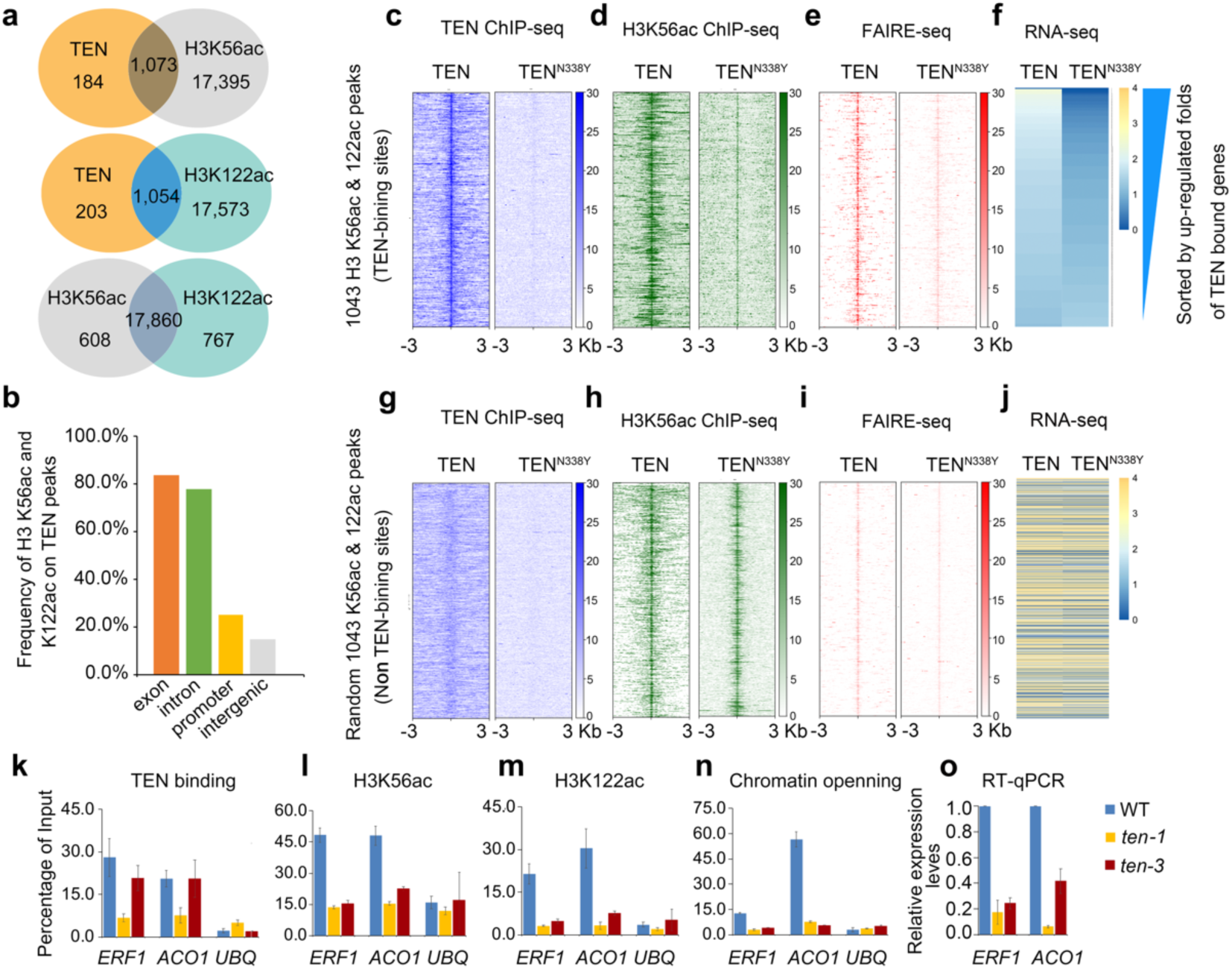
TEN promotes chromatin accessibility. **a**, Metagene analysis showing genome-wide colocalization of TEN intragenic peaks with H3K56ac and H3K122ac in tendrils. **b**, H3K56ac and H3K122ac are enriched in intragenic regions of TEN peaks, but not in promoter or intergenic regions. **c** to **f**, Read density heatmaps showing the intensity of TEN peaks (**c**), H3K56ac/H3 signals (**d**), chromatin opening signals (**e**) and RNA-seq signals (**f**) in WT and *ten-1* at 1043 overlapped peaks spanning ±3 kb from the center of the TEN peaks. Analyzed peaks were organized from top to bottom based on downregulation (fold) in *ten-1*. **g** to **i**, Read density heatmaps showing the intensity of TEN peaks (**g**), H3K56ac/H3 signals (**h**), FAIRE-seq signals (**i**) and RNA-seq signals (**j**) in WT and *ten-1* at 1043 random non-TEN binding peaks spanning ±3 kb from the center of the H3K56ac peaks. **k** to **o**, Validation of TEN binding, histone acetylation, chromatin opening and gene expression. Association of TEN protein with *ACO1, ERF1* loci and control regions (*UBQ, Csa3G778350*). ChIP was performed on WT, *ten-1* and *ten-3* with a TEN polyclonal antibody (**k**). H3K56ac (**l**) and H3K122ac (**m**) levels at the *ACO1, ERF1* and *UBQ* loci in WT, *ten-1* and *ten-3*. Chromatin opening detected at *ACO1, ERF1* and *UBQ* loci in WT, *ten-1* and *ten-3* by FAIRE-qPCR (**n**). Relative mRNA expression levels of *ACO1* and *ERF1* genes among WT, *ten-1* and *ten-3* detected by RT-qPCR, *UBQ* was used as internal control (mean ± SEM, *n* = 3) (**o**).

To assess the *in vivo* effect of the TEN C-terminus on TEN binding and HAT activities, we next examined the 1043 intragenic TEN binding sites that were enriched simultaneously for H3K56ac and H3K122ac in WT versus *ten-1* (Fig. 5c-f), as well as 1043 randomly selected H3K56ac and H3K122ac peaks lacking TEN binding as an internal control (Fig. 5g-j). TEN^N338Y^ lost most of its binding peaks in the *ten-1* mutant (Fig. 5c), consistent with our *in vitro* binding assays (Fig. 2o). This reduction of TEN binding, in the *ten-1* mutant, was associated with a decrease in H3K56ac levels (Fig. 5d). As H3K56ac and H3K122ac are implicated in the loosening and eviction of nucleosomes^20,26-28^, we reasoned that TEN may promote chromatin accessibility and, thereby, stimulate transcription of its target genes.

To test this notion, chromatin accessibility was tracked in the tendril genome by using FAIRE-seq (formaldehyde-assisted isolation of regulatory elements sequencing). Here, a strong correlation was observed between TEN binding, histone acetylation and chromatin accessibility (Fig. 5c-e); these functions were impeded dramatically in *ten-1* compared to WT (Fig. 5e). Importantly, RNA-seq analysis further showed that most genes that corresponded to the TEN intragenic peaks were downregulated in *ten-1* (Fig. 5f and Supplementary Table 6). By contrast, the internal controls exhibited no significant difference between WT and *ten-1* (Fig. 5g-j). The genomic view of two exemplary targets, *ACO1* and *ERF1*, also supported these findings (Supplementary Fig. 7c-d).

To survey the *in vivo* effect of the N121 domain on TEN binding, H3K56ac and H3K122ac, chromatin accessibility, and target gene expression, ChIP-qPCR, FAIRE-qPCR and qRT-PCR were performed at the TEN target genes, *ACO1* and *ERF1*, in WT and *ten-3* that has an in-frame deletion of Asp^55^ in the N121 domain, using *ten-1* as a negative control (Fig. 5k-o). As expected, at their TEN binding regions, both genes showed significant decreases in TEN binding (Fig. 5k), H3K56ac and H3K122ac levels (Fig. 5l-m, respectively) and FAIRE signals (Fig. 5n), as well as substantially reduced expression, being about 7-fold for *ERF1* and 17-fold for *ACO1* in the *ten-1* mutant (Fig. 5o).

Furthermore, we established that deletion of Asp^55^ in *ten-3* did not affect significantly the TEN binding levels (Fig. 5k). However, H3K56ac and H3K122ac levels, as well as chromatin accessibility, were significantly reduced (Fig. 5l-n). As predicted, these two genes were also downregulated by about 5-fold for *ERF1* and 3-fold for *ACO1* compared to WT (Fig. 5o), consistent with our biochemical result that Asp^55^ contributed largely to the HAT activity of N121 (Fig. 4h-i). These findings provide strong support for the hypothesis that, in tendrils, TEN binds specifically to its target intragenic regions, where it acetylates the globular domain of histone H3 to facilitate chromatin accessibility and, thereby activates its target gene expression.

Of equal importantly, combined with the phenotypic changes of tendril induced by the mutations in both the C- and N-termini, our findings provide *in planta* evidence that TEN binding, and the associated acetylation of the globular domain of H3, are critical for normal tendril architecture and behavior.

### Conserved HAT function of CYC/TB1 TFs conferred by IDR

To explore whether the N terminal regions of CYC/TB1-like proteins have conserved HAT activity, we performed N-terminal alignments of several CYC/TB1 TFs from among angiosperm species (Supplementary Fig. 8a), and cucumber TEN, durio CYC, Arabidopsis BRC1and BRC2, and maize TB1 were selected for further analysis. All these N-terminal regions contain a large portion of IDRs (Fig. 6a and Supplementary Fig. 8b) and share little sequence homology (Fig. 6b). However, these regions did contain some amino acids, including glutamine (Gln^14^), aspartate (Asp^55^), and cysteine (Cys^104^) that are responsible for the catalytic activity of N121 (Figs. 6b and 4h). As IDRs are generally evolving rapidly, at the primary sequence level, it is currently difficult to predict whether their functional consequences are preserved during the evolution of CYC/TB1-like proteins^45^.

**Figure 6.**
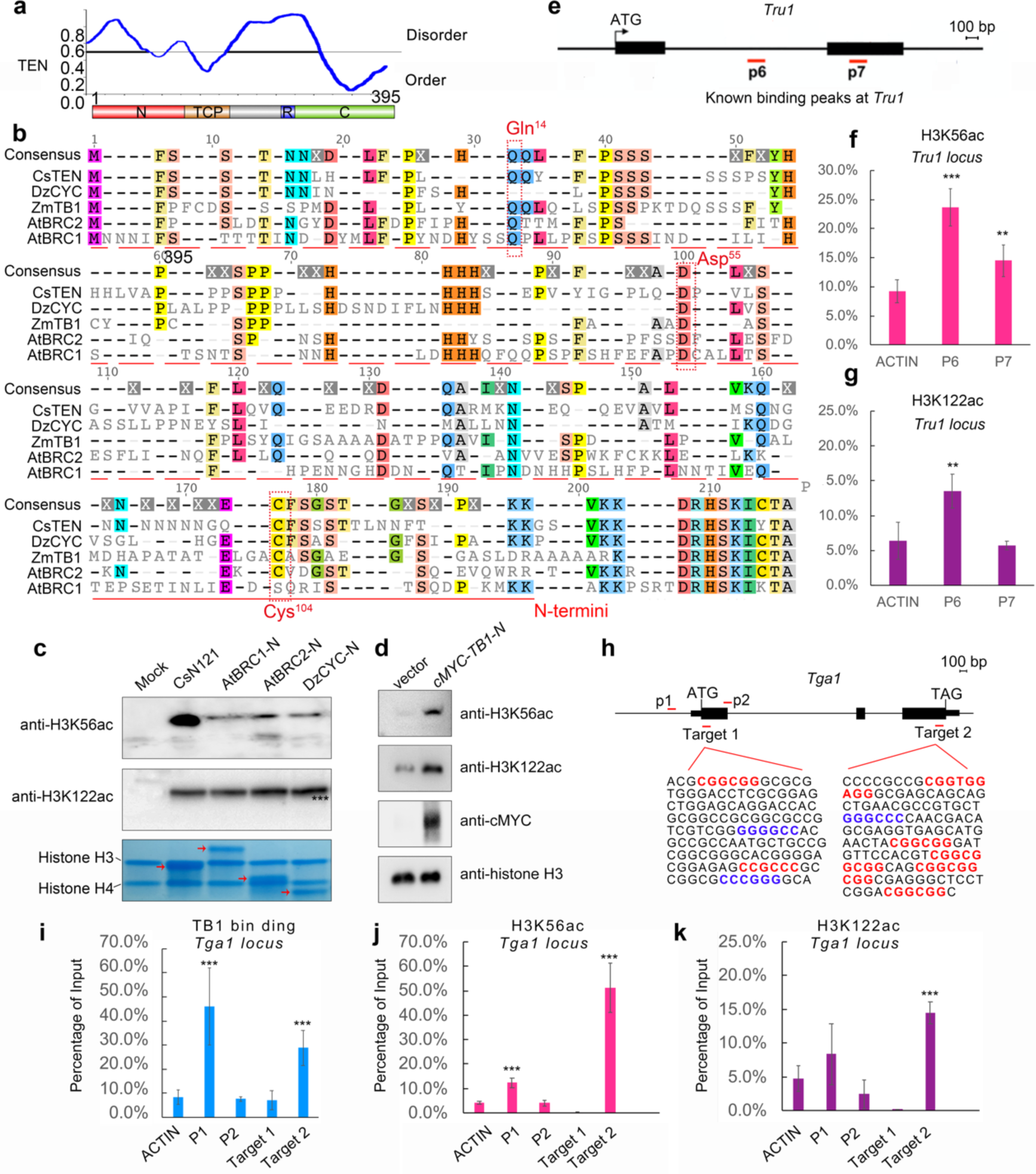
The N termini of CYC/TB1-like proteins have conserved HAT activities. **a**, IDRs of TEN. Graphs plotting intrinsic disorder (PONDR VL3-BA) for TEN protein. PONDR VL3-BA score (y-axis) and amino acid position (x-axis) are shown, indicating the intrinsically disordered and ordered regions. **b**, Alignment of the N-terminus of CYC/TB1-like proteins among four angiosperm species. Red dashed boxes indicate the Gln^14^, Asp^55^ and Cys^104^ amino acid residues. **c**, Acetylation of H3K56 and K122 within the H3-H4 tetramer by N terminus proteins of TEN, BRC1, BRC2 and durian CYC, determined by immunoblotting analysis. **d**, Overexpression of c-Myc-ZmTB1-N in tobacco leaves increased acetylation levels of H3K56 and H3K122. **e** and **f**, H3K56ac (**e**) and H3K122ac (**f**) levels at P6 and P7 sites of *Tru1* locus bound by TB1. **g**, Diagram of the *Tga1* genic region. Black boxes indicate exons, and lines between boxes represent introns. Locations of the amplicons (targets 1 and 2) used for ChIP-qPCR are marked below. Potential TB1-binding motifs are highlighted in blue (GGGCCC) and red (CCNCCN). **h** to **j**, TB1 binding (**h**), H3K56ac (**i**) and H3K122ac (**j**) levels at target 1 and 2 sites of *Tga1*. ChIP was performed with antibodies to TB1, H3K56ac and H3K122ac (mean ± SEM, *n* = 3). Triple asterisk, *P* <0.01.

From the view of this situation, N terminal proteins of DzCYC, AtBRC1 and AtBRC2 were expressed and purified to test their effects on H3K56 and K122 acetylation (Supplementary Fig. 9). Our results showed that each protein acetylated both K56 and K122 in the histone H3 globular domain (Fig. 6c and Supplementary Fig. 10). Transient expression of ZmTB1 in *Nicotiana tabacum* leaves similarly resulted in increased H3K56 and H3K122 acetylation (Fig. 6d). These findings support the hypothesis that such homologous IDRs retain similar functions, despite extensive sequence divergence^45^.

Recent studies showed that ZmTB1 bound to the gene bodies of a target gene, *Tru1*^7^. The acetylation levels of TB1-binding sites on *Tru1*, in maize tiller buds, were assessed by ChIP-qPCR, using the primers previously reported (Fig. 6e)^7^. Both H3K56ac and H3K122ac levels were enriched significantly at the intronic P6 site (Fig. 6f-g), consistent with previous results that TB1 binds to the P6 site with high affinity^7^. *Teosinte glume architecture1* (*Tga1*) is another maize domestication gene that is downstream of TB1 and upregulated by TB1^46^. By using a TB1 antibody, we established that, in addition to binding to the known p1 site in the promoter, TB1 could bind to a DNA element located in the third exon of the *Tga1* locus (Fig. 6h-i). Furthermore, we showed that both H3K56ac and H3K122ac modifications were enriched significantly at the TB1 intragenic binding site (Fig. 6j-k). H3K56ac was also identified to be significantly enriched at the p1 promoter region bound by TB1. However, although TB1 showed stronger binding at the promoter, the H3K56ac level was lower than that in the intragenic binding site (Fig. 6i-j). We inferred that this might be attributable to acetylation at the histone H3 globular domain, such as H3K56ac or K122ac in coding regions, thereby evading the surveillance of the classic histone deacetylation pathway^14^. Recent studies indicated that AtBRC1 also bound to the introns of their targets *HB53 and HB21*^8^, and activated their expression. Our study implicates a conserved mechanism for gene activation, by CYC/TB1-like TFs, which function both as TFs binding at intragenic enhancer sites, and as HATs acting on the histone globular domain (Supplementary Fig. 11). This discovery provides important insights into the regulatory mechanism involved in axillary bud development and, thus, plant architecture.

## DISCUSSION

Each genome has undergone a unique evolutionary trajectory, offering a distinct window to explore a unique range of biological processes. In this study, the cucumber genome offered a special lens to observe the molecular functions of the CYC/TB1-like protein family of TFs, to gain molecular insight into how intragenic binding, by a TF, can regulate gene expression, to explore the function of IDR within TFs.

A recent genome-wide ChIP-seq assay reported that TB1 mainly binds to promoters, and only a few peaks were located within gene body regions^6^. In our study, TEN predominantly bound to the gene body, and this was not unexpected, given that these two genes have diverged over a significant period of evolution. In addition, as ∼1 mm-long arrested buds were used for TB1 ChIP-seq, whereas tendrils at the coiling stage were used for TEN ChIP-seq assays in the current study, we cannot preclude the possibility that TEN may have another binding specificity at the stage of tendril meristem development. We further compared the regulatory datasets of TB1 and TEN. Despite the quite different genome-wide binding features, they also exhibited a number of similar genetic pathways involved in regulating phytohormones and trehalose 6-phosphate metabolism (Supplementary Fig.12 and Supplementary Table 7).

In this study, we discovered three features of TEN; i.e., 1) its function may primarily be as a transcriptional activator, 2) it can bind intragenic enhancers of target genes, and 3) it is a novel, non-canonical HAT that acts on the histone globular domain. TEN, itself, then becomes the link that connects these three activities, and therefore, provides an answer to the question regarding the mechanism by which the modification at the H3 globular domain is integrated into the transcriptional process^32^. Tail-based histone acetylation sites function as platforms for the recruitment of transcriptional regulators, which usually occurs around transcription start sites. Our study supports the notion that acetylation of the histone globular domain, in concert with the maintenance of accessible chromatin, are important facets of transcriptional regulation, via intragenic enhancers. In view of these findings, we investigated the genomic distribution of two human enhancer binding TFs, the heat shock factor (HSF-1)^47^ and the estrogen receptor alpha (ERα)^20,48^, both of which can interact and recruit CBP/p300. Importantly, we showed that H3K56ac or H3K122ac modification was observed to more likely occur on exons than the other regions bound by the HSF-1 or ERα (Supplementary Fig. 13a-b). Utilizing K56ac or K122ac could also be advantageous, as it can evade the surveillance of the classic histone deacetylation pathway in coding regions^14^. Thus, our findings add critical information that advances knowledge as to how the binding of a TF, within the intragenic region of a gene, can influence expression. The acetylation and loosening of chromatin, at specific intragenic targets, by TEN, could well act to facilitate productive RNA elongation by RNA polymerase (supplementary Fig. 11). Besides, it has not escaped our notice that the TEN-dependent epigenetic regulation, on these intragenic targets, might also direct codon choice and affect protein evolution^49^ during the evolutionary process of tendril formation.

HATs form a diverse collection of enzymes characterized by their sequence homology and structural features, including the GNAT, MYST and p300/CBP families^50^. These classical HATs are mainly recruited to target promoters through physical interactions with sequence-specific TFs^51^, whereas in mammalian systems, some HATs also possess DNA binding activity^52,53^. In our study, we discovered a non-canonical type of DNA binding HAT – TEN, in plants, that acetylates the histone H3 globular domain at intragenic enhancers, through its N terminus which harbors a significant portion of IDRs (Fig. 6a and Supplementary Fig. 8). We also showed that the kinetic of acetylation capacity of N121, on H3K56 and H3K122, is comparable to that of Rtt109, and is approx. one-eighth of P300. For Rtt109, studies have demonstrated that Vps75 greatly enhances its HAT activity^44^, and therefore, the possibility exists that TEN may similarly have increased HAT activity, *in vivo*, through its interaction with other proteins, or via posttranslational modifications.

Due to the highly diverged primary sequences, the function and evolution of these IDRs has remained largely unexplored. Our findings, that the intrinsically disordered N-termini of all tested CYC/TB1-like proteins have conserved HAT activity, now provide an answer to the frontier question regarding the mechanism of action of TFs with IDRs^35^. In addition, recently, IDRs were shown to play an important role in the compartmentalization of the transcription apparatus, and the formation of liquid-liquid phase separation, related to the nature of super-enhancers^13^. In this regard, our study also offers a possibility to test if the typical intragenic enhancer works by a similar mechanism.

## Acknowledgments

We thank Yihua Huang and John Doebley for comments on the manuscript, Jinsheng Lai, Kehui Liu, Haiteng Deng, Shilong Fan, Xiling Chen, Lvjun Guo and Yu Zhao for experimental assistance, and Huai Wang for providing TB1 antibodies.

## Author Contributions

S.H. and X.Y. designed the research. X.Y., Z.Z., and B.W. made major contributions to biochemical analyses and ChIP assays. J.Y. contributed to protein purification. T.L., X.Y. and Z.Z. led bioinformatic analyses. X.Y. and T.X. led genetic transformation of plants. S.W. helped to collect tendril materials. G.L., J.Z., and Z.Z. contributed to histone purification and assembly. S.H., X.Y., G.L., W.J.L., and J.Y. analyzed the data and wrote the manuscript.

## Methods

### Experimental materials

The cucumber (*Cucumis sativus* L.) inbred line 404 (WT) and its BC_3_S_2_ mutant tendril near-isogenic line 404-38 (*ten*) were used for genome-wide ChIP-seq, FAIRE-seq and RNA-seq analyses. After seed germination, 100 plants of each line were grown, in a greenhouse, in pots containing mixed peat moss and vermiculite (v/v = 1:1) and were transplanted to soil at the three-leaf stage. Pest control was performed according to standard management practices.

The cucumber inbred line CU2 was used in cucumber transformation. Seeds were soaked in distilled water, at 50°C for 30 min. Seed coats were removed, and the naked seed was then surface-sterilized by sequential immersion in 70% ethanol for 15 s and 0.6% sodium hypochlorite solution for 15 min, followed by eight rinses in sterile distilled water. Sterilized seeds were spread on 1× Murashige and Skoog medium (Phytotech, Cat. #M519), supplemented with 2 mg/L 6-Benzylaminopurine (Sigma, Cat. #B3408) and 1 mg/L ABA (Phytotech, Cat. #A102), for two days at 28°C. Cotyledons were excised from germinated seedlings and infected with Agrobacterium. Subsequently, after shoot regeneration, elongation, and rooting processes, the rooted plants were transplanted to the greenhouse.

For transient expression analysis, tobacco plants (*Nicotiana benthamiana*) were grown in pots containing mixed peat moss and vermiculite (v/v = 1:1) in a growth chamber with a light regime of 16 h light/8 h dark at 22°C.

### Plasmid construction and plant transformation

To generate CRISPR/Cas9 engineered mutations in the *TEN* gene, a binary CRIPSR/Cas9 vector pBSE402 plus a *35S-GFP* expression cassette was modified from pBSE401a^54^. For assembly of *TEN* sgRNA into pBSE402, equal volumes of 100 μM forward and reverse primers were mixed, incubated at 95°C for 5 min, and slowly cooled to room temperature, resulting in a double stranded DNA fragment with sticky BsaI ends. This short DNA fragment was then assembled into pBSE402, by restriction fragment ligation, using BsaI and T4 Ligase (New England Biolabs). Primers are shown in Supplementary Table 8. *Agrobacterium tumefacines* strain EH105, carrying a pBSE402-TEN construct, was used to transform the cucumber inbred line CU2, using cotyledonary nodes as explants, as previously described^55^. Shoot regeneration, elongation and rooting processes strictly followed normative procedures.

Genomic DNA was extracted from the positive transgenic plants using the DNeasy Plant Mini Kit (Qiagen, Cat. #69104). PCR was performed using gene-specific primers (Supplementary Table 8). PCR products were cloned into pEASY-Blunt Zero (TRANSGEN BIOTECH, Cat. #CB501) and the various alleles for the *TEN* gene were identified by sequencing.

### ChIP-seq and ChIP-qPCR

ChIP assays were performed, as described previously^56^, with some modifications. Briefly, normal tendrils (WT), at coiling stage (the status at which the tendril attaches to a support, but before free tendril coiling occurs), mutant tendrils of *ten-3* (at the corresponding growth status where the tendril would normally be attaching to a support) and *ten-1* mutants (at the corresponding growth status where the modified tendrils show slight curling of petioles) (Fig. 1g) were used in ChIP assays for two biological replicates. Harvested tendrils (30 g of each material divided into 10 equal samples) were fixed in cross-linking buffer (10 mM sodium phosphate, pH 7.0, 50 mM NaCl, 0.1 M sucrose, and 1% formaldehyde) under vacuum for 10 min. Fixation was stopped by incubation in 0.25 M glycine for an additional 10 min. Tendril material was then ground, in liquid nitrogen, and 3 g aliquots of powdered tissue were resuspended in 30 mL of extraction buffer (0.4 M sucrose, 10 mM Tris-HCl pH 8.0, 5 mM β-mercaptoethanol, 1 mM PMSF, and Protease Inhibitor cocktail). Chromatin isolation was performed, as described previously^56^, and then sonicated (Ningbo Scientz Biotechnology, JY96-IIN) for 12 cycles, each with a 15 s pulse at 70% of maximal power, followed by a 45 s cooling period, on ice, to achieve an average DNA size of 200 bp for immunoprecipitation. The following antibodies were used for ChIP assays: TEN antibody, anti-H3K56ac (active motif, Cat. #39282) and anit-H3K122ac (Abcam, Cat. #Ab33309) (Supplementary Table 9). ChIP products were combined and eluted into 50 μL of TE buffer for ChIP-seq (5 ng DNA) or ChIP qPCR (1 μL aliquot).

ChIP-qPCR was performed, as described previously^56^. Primers are listed in Supplementary Table 8, and *UBQ* (*Csa3G778350*), the ubiquitin gene, was used as a negative control. The qPCR signals derived from the ChIP samples were normalized to the signals derived from the input DNA control sample. The value (percentage of input; input %) was calculated by the 2^-ΔCt^ method.

### FAIRE

FAIRE assays were performed, as described previously^57^. Normal tendrils (WT), at the coiling stage, and mutant tendrils (*ten-3* and *ten-1* mutants), at the corresponding growth stage, were used in FAIRE assays. Two grams of tissue were fixed with formaldehyde and regulatory elements were isolated. FAIRE DNA was dissolved into 30 μL of TE buffer for FAIRE-seq (5 ng DNA) or FAIRE qPCR (1 μL aliquot). Two biological replicates were performed for each FAIRE assay.

qPCR was performed on crosslinked and non-crosslinked (input) FAIRE samples. The ubiquitin gene, *UBQ*, was used as a negative control. Lists of all primers used are given in Supplementary Table 8. The value for DNA accessibility over that of input was obtained by the 2^-ΔCt^ method.

### RT-qPCR analyses

Total RNA was isolated using an RNA extraction Kit (Qiagen, Cat. #74903). First-strand cDNA was synthesized from 1 μg total RNA using the M-MLV Reverse Transcriptase (Promega, Cat. #M1705) (primers are listed in Supplementary Table 8). Primer specificity was checked by sequencing and blast analysis. qPCRs were performed on an ABI 7900, using SYBR Premix (Roche, Cat. #4913914001), according to the manufacturer’s instructions. Three technical replicates and three independent biological experiments were performed in all cases. Relative gene expression was assessed using the comparative 2^-ΔΔCt^ method. *UBQ* was used as an internal reference gene.

### High-throughput sequencing

ChIP-seq or FAIRE-seq libraries were prepared using the Illumina ChIP-seq DNA Sample Prep kit, according to manufacturer’s instructions, with the following modifications: mRNA adaptor indexes from the TruSeq RNA kit were used, and enrichment PCR was also performed with reagents from the Illumina TruSeq mRNA kit. The enriched libraries were purified, with AMPure magnetic beads (Agencourt), the concentrations checked with Qubit (Invitrogen), and the distribution and size of fragments were confirmed with a Bioanalyzer (Agilent). Four samples were pooled in equimolar quantity and sequenced on a HiSeq2500 (single read, 50 bp) to yield up to 30 million reads per sample. The obtained reads were demultiplexed with Illumina CASAVA 1.8 software.

RNA-seq libraries were developed from five biological replicates from tendrils (WT) and mutant tendrils (*ten*). The 100 bp paired-end reads (2.4 Gb, 10×) for each sample were generated from the RNA-seq libraries with an Illumina HiSeq 2500 sequencer.

### Mapping of sequencing reads and data analysis

All sequencing reads were mapped to the cucumber genome, using bowtie software (http://bowtie-bio.sourceforge.net)^58^, with default parameters, except for discarding multiple loci-matching reads that might introduce error signals by repeat counting.

For computational processing of ChIP-seq: Peaks for TEN and TEN^N338Y^ were identified by the model-based analysis software MACS (http://liulab.dfci.harvard.Edu/MACS/)^59^, using input DNA as a control. MACS default parameters were used, except for detecting more reliable TEN association signals with –mfold = 0, 30 and fold enrichment ≥ 2. Peaks for H3K56ac and H3K122ac were identified by RSEG software (https://github.com/smithlabcode/rseg)^60^, under default parameters, using input DNA as a control. For computational processing of FAIRE-seq data: Data peaks were identified by F-Seq software (http://fureylab.web.unc.edu/software/fseq/)^61^, using input DNA as a control. Heatmap graphs of peaks were plotted using deeptools software (https://github.com/deeptools/deepTools) with normalization to 1×.

For computational processing, RNA-seq data were mapped to the cucumber genome, using tophat2 software (http://tophat.cbcb.umd.edu/)^62^, with default parameters. According to the cucumber genome annotation, all mapped reads were then assembled into known transcripts by Cufflink software. Next, the expression of transcripts was calculated in fragments per kilobase of exon model per million mapped fragments.

The putative DNA-binding motifs in the TEN binding peaks were searched using MEME-ChIP software^63^ (the online version 5.0.5), by using the default background model of MEME-ChIP. The background model is normalized for biased distribution of letters in the input sequences.

### Purification of TEN protein from Sf9 insect cells and *E.coli*

The cDNA of *TEN, TEN*^*N338Y*^, *N121* and *ΔN121* was cloned into pFast-FH vector (inserting a FLAG tag into pFast-HTB, Life Technologies, Cat. #10712-024), respectively. Primers for constructing these vectors are shown in Supplementary Table 8. Bacmid preparation and insect cell transfection were conducted using the Bac-to-Bac® Baculovirus Expression System, according to the manufacturer’s instructions. The isolated P3 recombinant baculoviruses were added to the cultured Sf9 cells, at a volume ratio of 1:100. Cells were collected after another 48–60 h of cultivation at 27°C and 110 rpm in Nalgene conical flasks.

Insect cells that expressed recombinant proteins were resuspended and sonicated in lysis buffer (50 mM HEPES-KOH, pH 7.5, 500 mM NaCl, 5% glycerol). Cell lysates were centrifuged at 16,000 × g for 60 min at 4°C. The supernatants from cell lysates were loaded onto a gravity column (Bio-Rad, Cat. #732-1010) filled with 2 mL Anti-FLAG M2 Affinity Gel (Sigma, Cat. #A2220), and bound proteins were eluted with a FLAG elution buffer (50 mM HEPES-KOH, pH 7.5, 500 mM NaCl, 5% glycerol, 0.4 mg/mL 1× FLAG peptide). The eluate was concentrated to 2 mL and then loaded onto a Superdex 200 column (GE Healthcare, Cat. #10034543) for size exclusion chromatography (S200 buffer: 50 mM Hepes-KOH, pH 7.5, 500 mM NaCl, 5% glycerol, 1 mM DTT). The fractions containing target protein were collected and constituted the purified protein.

The cDNA of a series of N-terminal sequences, including the N121, the point mutated N121 and N-termini of the TB1 TF family, were cloned into the pET22b vector, fused to the N-terminus with a 6× His-tag. The resultant plasmid was transformed to *E. coli* BL21(DE3), then identified by PCR, double enzyme digesting (*Ned* I and *Xho* I) and sequenced. Expression of the recombinant N121 protein was induced, with 0.2 mM isopropyl thiogalactoside for 5 h at 26°C, and then affinity purified, using lysis buffer (25 mM Tris-HCl, pH 7.5, 500 mM NaCl), in combination with an Ni^2+^-chelating Sepharose Fast Flow (Amersham Biosciences) column, following the manufacturer’s instructions.

### Electrophoretic mobility-shift assays (EMSA)

EMSA was performed using recombinant MBP-TCP, MBP-TCP+R, FLAG-ΔN121, FLAG-TEN, or FLAG-TEN^N338Y^ protein purified from insect cells. DNA probes containing the CTCCGCC motif, mutant CTAAGCC motif, or a GTGGTCCCAC motif, used for the EMSA, were synthesized and amplified, by PCR, using the biotin-labeled primers listed in Supplementary Table 8. Binding reactions were performed in 20 μL of binding buffer, composed of 10 mM Tris-HCl, pH 7.5, 200 mM NaCl, 10 mM KCl, 1 mM MgCl2, 10 μM ZnCl2, 0.5 mg/mL BSA, 0.02 mg/mL of poly (deoxyinosinic-deoxycytidylic) sodium salt [poly (dI-dC)] (Thermo Scientific, Cat. #20148E), 1 mM DTT and 10% glycerol. Binding reactions were carried out using 0, 100 or 200 ng of recombinant protein and 5 nM of each biotin-labeled probe, at 4°C for 1 h. Samples were separated on 6% polyacrylamide gels (19:1 acryl:bisacrylamide) in Tris-borate-EDTA, at 4°C. After the sample transfer, the PVDF membrane was exposed under ultraviolet light to cross-link the samples, and then blocked with blocking reagent (provided in the kit) for 15 min. Biotin signal was visualized using a Chemiluminescent Nucleic Acid Detection Module kit (Thermo Scientific, Cat. #89880). Competition experiments were performed using from 100- or 1000-fold levels of unlabeled fragments.

### Expression and purification of recombinant histones

Recombinant histones were expressed in BL21 (DE3) pLysS. Single colonies were grown in 1 L of lysogeny broth (LB) medium at 37 °C until reaching an optical density at 600 nm (OD600) = 0.6, and induced with 0.5 mM IPTG for 2 h. Cells were harvested by centrifugation, at 5,000 × g, and resuspended in lysis buffer (50 mM Tris, 100 mM NaCl, 1 mM EDTA, 1 mM 2-mercaptoethanol, pH 7.5). Cells were lysed, by sonication, and then centrifuged at 30,000 × g.

For histone purification, inclusion bodies from 1L bacterial culture were washed by resuspension in 100 mL wash buffer, plus 1% Triton X-100, and then centrifuged for 10 min at 4°C at 23,000 × g. This step was repeated, once, with wash buffer plus Triton X-100, and twice with wash buffer. The pellet was solubilized in 30 mL unfolding buffer (7 M guanidium hydrochloride, 20 mM Tris, pH 7.5, 10 mM DTT) for 1 h, at RT. After centrifugation the supernatant was analyzed by SDS-PAGE.

### Histone tetramer and octamer reconstitution

For histone H3-H4 tetramer or histone octamer reconstitution, histone H3, H4 or H2A, H2B, H3 and H4 were mix, at equimolar amounts, and then dialyzed at 4°C against three changes of 2 L freshly-prepared refolding buffer (2 M NaCl, 10 mM Tris-HCl, pH 7.5, 1 mM EDTA, 5 mM β-mercaptoethanol) for 12 h. Next, the mixture was dialyzed for another 12 h, in 2 L fresh refolding buffer. After centrifugation, the supernatant was collected and concentrated to a final volume of 500 μL. Following a minimum of three centrifugation steps, the cleared supernatant was loaded onto a Superdex200 gel filtration column and equilibrated with refolding buffer. The purity and stoichiometry of eluted fractions were checked by SDS-PAGE. Histone H3-H4 tetramer or histone octamer peak fractions were collected, together, and store at -80°C.

### *In vitro* nucleosome assembly

Mononucleosomes were assembled on 208 bp (*5S* rDNA) DNA fragments. Before adding octamer, the salt concentration of the DNA solution was adjusted to 2 M, using 5 M NaCl and TE. DNA and histone octamers were mixed at a 1:1.05 molar ratio in 2 M NaCl buffer. A peristaltic pump was used for continuous dialysis against 450 mL of refolding buffer (2 M NaCl, 10 mM Tris-HCl, pH 7.5, 1 mM EDTA, 5 mM β-mercaptoethanol) for 16 h at 4°C, under constant stirring, with continuous addition of TE buffer, into the dialysis buffer, to reduce the salt concentration to 0.6 M. Samples were collected after final dialysis in HE buffer (10 mM Hepes, pH 8.0, 0.1 mM EDTA) for 4 h. The assembled nucleosomes were visualized on 2% agarose gels.

### Histone acetyltransferase assay

HAT assays were performed in a 30 μL reaction medium, using either 0.5 μg of recombinant histone H3 (Millipore, Cat. #14-411), 2 μg of chicken core histones (Millipore, Cat. #14-411), 1 μg H3-H4 tetramer, 2 μg histone octamer or 2 μg mononucleosome, in the presence of 1 μCi of ^3^H-acetyl-CoA (ARC, Cat. #0213A-50 µCi) or 200 μM acetyl-CoA. Enzymatic reactions were performed using 100 nM of purified FLAG-N121, and the same amount of FLAG-TEN, FLAG-TEN^N338Y^, FLAG-ΔN121 or commercial P300 protein. Reactions were incubated at 30°C for 2 h, and then 7.5 μL 5× SDS-PAGE sample buffer was added, followed by boiling for 5 min. After being resolved on 15% SDS-PAGE, aliquots were subjected to LC-MS/MS, autoradiography or immunoblotting. For autoradiography, proteins were separated and transferred to a PVDF membrane, using a semi-dry blotter (TE70, GE Life Sciences). ^3^H signal was detected with a BioMax Transcreen Intensifying Screen LE (Sigma, Cat. #Z374318) and BIOMAX MS films. For immunoblotting, the antibodies specific for different acetylated lysine residues were listed in Supplementary Table 9.

### LC-MS/MS analysis

Equal protein amounts were separated by SDS-PAGE. The gel bands of histone H3 protein were excised, reduced with 25 mM of DTT and alkylated with 55 mM iodoacetamide, followed by addition of propionic anhydride and in-gel digestion, overnight, with sequencing-grade modified trypsin, at 37°C. Peptides were extracted twice with 0.1% trifluoroacetic acid in 50% acetonitrile aqueous solution for 30 min and then dried in a speedvac. Peptides were redissolved in 25 μL 0.1% trifluoroacetic acid and 6 μL of extracted peptides were analyzed by Q Exactive HF-X mass spectrometer.

In LC-MS/MS analysis, digestion products were separated by a 120 min gradient elution, at a flow rate of 0.300 µL/min, using a Dionex 3000 nano-HPLC system, which was directly interfaced with a Thermo Q Exactive HF-X mass spectrometer. The analytical column was a fused silica capillary column (75 µm ID, 150 mm length; packed with C-18 resin). Mobile phase A consisted of 0.1% formic acid, and mobile phase B consisted of 80% acetonitrile and 0.08% formic acid. The Q Exactive mass spectrometer was operated in the data-dependent acquisition mode, using Xcalibur4.1 software, and a single full-scan mass spectrum in the Orbitrap (300-1800 *m/z*, 12,000 resolution) was followed by 40 data-dependent MS/MS scans. The MS/MS spectra from each LC-MS/MS run were searched against the selected database, using Proteome Discovery searching algorithm (version 1.4).

The MS/MS spectra from each LC-MS/MS run were searched against the Histone H3.fasta file. The search criteria were as follows: full tryptic specificity was required; two missed cleavages were allowed; carbamidomethylation© were set as the fixed modification; the oxidation (M), propionyl (P) and acetyl (K) were set as the variable modification; precursor ion mass tolerances were set at 20 ppm for all MS acquired in the orbitrap mass analyzer; and the fragment ion mass tolerance was set at 0.02 Da for all MS2 spectra acquired. The peptide false discovery rate (FDR) was calculated using Percolator provided by PD. When the q value was smaller than 1%, the peptide spectrum match (PSM) was considered to be correct. FDR was determined based on PSMs when searched against the reverse, decoy database. Peptides assigned only to a given protein group were considered as unique. The FDR was also set to 0.01 for protein identifications. The peak areas of fragment ions were used to calculate the relative intensity of precursor ion for selected peptides.

### Quantitative calculations for residue-specific histone acetylation

The method for quantitative calculation was performed, as described previously^41^. The fraction of a specific peptide (F_s_) was calculated by Equation 1, where I_s_ was the intensity of an acetylated peptide state, and I_p_ was the intensity of any state of that peptide.

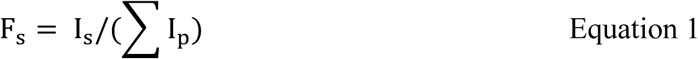

The concentration of acetylation at specific lysine residues was quantified and calculated by multiplying the fraction F_s_ by the initial concentration of histone.

For steady-state kinetic analyses, all models were fitted to the data, using Prism (version 7.0). The initial rates (V) of acetylation were calculated from the linear stage in acetylation for a 10 min reaction time. To measure steady-state parameters for H3-H4 tetramer, *K*_cat_ and *K*_m_ were determined based on the equation:

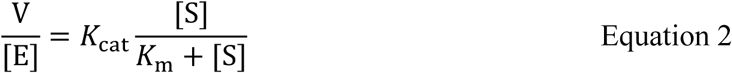

where [S] was the concentration of substrate (H3-H4 tetramer), [E] was the concentration of N121 protein, and V was the initial rate of acetylation.

### Protein extraction and detection by immunoblotting

The 18-residue peptide, LNNFTKKGSVKKDRHSKC, spanning the N121 and TCP domains, was selected as antigen for polyclonal antibody production. Two grams of cucumber tendrils, or tobacco leaves, were harvested and ground to fine power in liquid N2. Total protein was extracted using a protein extraction buffer (20 mM Tris-HCl, pH 7.5, 150 mM NaCl, 4 M Urea, 10% glycerol, 5 mM DTT, 1 mM PMSF and 1× protease inhibitor cocktail). The samples were centrifuged at 12,000 × g at 4°C for 30 min. Total protein was quantified using BCA Protein Assay Kit, according to the manufacturer’s instructions. And 20–50 μg protein sample was resolved on 12.5% SDS-PAGE and then transferred to a PVDF membrane, using a semi-dry blotter. Western blot analyses were preformed using antibodies listed in Supplementary Table 9. Immunoblotting signal was visualized using a SuperSignal West Femto kit (Thermo Scientific).

### Immunolabelling

Half of each tobacco leaf was infiltrated with *A. tumefaciens* strains GV3101 expressing c-Myc-TEN, and the other half without infiltration was used as a control. After 72 h post infiltration, the entire tobacco leaf was cut into 0.5 cm × 1 cm pieces and fixed in cold 4% paraformaldehyde in Tris-HCl buffer (10 mM Tris, pH 7.5, 100 mM NaCl, 10 mM EDTA) for 20 min. Leaf pieces were then washed, twice, using ice-cold Tris-HCl buffer for 10 min each and nuclei were released by finely chopping in LB01 buffer (15 mM Tris-HCl, pH 7.5, 2 mM EDTA, 0.5 mM spermine, 80 mM KCl, 20 mM NaCl, 0.1% Triton X-100), followed by filtration through a cell strainer cup (BD falcon). Nuclei in the flow-through were then 1:4 diluted in sorting buffer (100 mM Tris, pH 7.5, 50 mM KCl, 2 mM MgCl_2_, 0.05% Tween 20, 5% sucrose), spotted onto microscopy slides, and air-dried. After post-fixation with 4% paraformaldehyde in PBS buffer (10 mM sodium phosphate, pH 7.0, 143 mM NaCl), slides were used for immunolabelling. Double labeling was performed using the c-Myc antibody (1:200), H3K56ac antibody (1:500) and H3K122ac antibody (1:500). C-Myc-TEN was detected by FITC-conjugated goat anti-rabbit (1:200, ZSGB-BIO) secondary antibodies, and each specific histone acetylation was visualized by TRITC-conjugated goat anti-rabbit (1:200, ZSGB-BIO) secondary antibodies. After staining, slides were mounted in mounting medium with DAPI and then photographed on a Leica TCS SP8 confocal microscope. More than fifty pairs of transfected nuclei versus non-transfected nuclei, in the same field of view, were observed to collect consistent results.

### Immunoprecipitation

Immunoprecipitation was performed to enrich the TEN protein for detection and LC-MS/MS analysis. 10 mg total tendril protein from the supernatants was incubated with an excess amount of anti-TEN antibody, at 4°C overnight with rotation, followed by addition of 50 μL Protein A Dynal beads (Thermo Scientific, Cat. #10002D) for an additional 2 h. Beads were then washed, 3 three times, with extraction buffer. Half of the immunoprecipitates was analyzed by immunoblotting with anti-TEN antibody, and the other half was resolved on 12.5% SDS-PAGE for LC-MS/MS analysis.

### Transient expression in tobacco leaves

TEN, TEN-N121, TEN-ΔN121 and TEN-ΔC130, fused with 5 × c-Myc tag peptides (EQKLISEEDL), were cloned into a binary vector (pCAMBIA1300) downstream of the *35S* promoter, using the primers listed in Supplementary Table 8. Constructs were transformed into *A. tumefaciens* strain GV3101. After cultivation, overnight, cells were harvested by centrifugation and resuspended in 10 mM MES (pH 5.6) buffer containing 10 mM MgCl_2_ and 200 μM acetosyringone (Sigma, Cat. #D134406) at OD600 = 1.0. After incubation, at room temperature for 3 h, in the dark, the *Agrobacterium* suspension was infiltrated into leaves of one-month-old tobacco plants from the adaxial side, using a needleless syringe. Leaf samples were harvested after 3 days and used for immunoblotting or immunolabelling analyses. These experiments were repeated, independently, at least three times with similar results.

### Data availability

Raw data were deposited at the Sequence Read Archive (SRA) under accession number PRJNA520931.

